# Robust footprinting with sample-specific Tn5 bias correction for bulk and single cell ATAC-seq

**DOI:** 10.1101/2025.10.17.683160

**Authors:** Yuxuan Lin, Hanzhi Wang, Parker C. Wilson, Nancy R. Zhang

## Abstract

Footprint analysis of assay for transposase-accessible chromatin via sequencing (ATAC-seq) enables base-resolution mapping of regulatory elements but is often underpowered and prone to false positives due to data sparsity and Tn5 transposase cleavage bias. We uncover substantial sample-to-sample variability in Tn5 transposase cleavage bias across samples, revealing a previously underappreciated source of batch effects and highlight the motivations for sample-specific Tn5 bias modeling. We present TraceBIND, a computational framework that corrects sample-specific Tn5 bias and use it to identify TF and nucleosome footprints through a dynamic flanking window statistical scan. Compared to existing approaches, TraceBIND substantially reduces false discoveries, controlling type 1 error while maintaining high sensitivity. Even though TraceBIND is completely unsupervised and does not require training on ChIP-seq data, we used multiple validation analyses to demonstrate that TraceBIND matches the power of supervised methods for detection of known TF binding sites. In scATAC-seq data from aging rat kidney, TraceBIND discovers age-associated dynamic regulatory changes, linking footprint activity to age-associated epigenetic drift. This demonstrates that footprint-informed scATAC-seq analysis reveals rich regulatory signals missed by conventional peak-based approaches.

## Introduction

A central goal in molecular biology is to understand how the regulatory genome controls gene expression across diverse cellular contexts. High-resolution maps of chromatin accessibility and regulatory element activity are essential for deciphering the regulatory network and linking it to cellular function and phenotype^1,2^. Chromatin immunoprecipitation followed by sequencing (ChIP-seq)^3^ has been employed for mapping regulatory proteins, but its requirement for factor-specific antibodies limits its scalability across all regulatory proteins and cellular contexts. By contrast, assays such as ATAC-seq have become indispensable for profiling chromatin state and inferring regulatory activity across the genome^4,5^. The recent advent of single-cell ATAC-seq (scATAC-seq)^6,7^ and single-cell multiome technologies^8–10^,which jointly measure chromatin accessibility and gene expression, has opened new frontiers in understanding gene regulation at cell-type resolution^11,12^. These developments have created a pressing need for robust analytical tools that can extract fine-scale regulatory insights from sparse and noisy chromatin accessibility data.

Among the most informative analyses of ATAC-seq data is footprinting, which aims to identify transcription factor (TF) binding and nucleosome positioning by detecting localized protection from transposase cleavage^13^. While conventional ATAC-seq analyses, especially in the single cell setting, rely on aggregate peak- or bin-level summaries^14,15^, these coarse features can miss the fine-scale binding signals that footprinting captures. Footprinting offers base-pair resolution and the potential to reveal dynamic gene regulatory activity, yet its application has been limited by technical challenges, particularly those arising from Tn5 cleavage bias^16^.

Tn5 cleavage is known to be sequence- and context-dependent^17^, introducing artifacts that can obscure true binding signals. Several methods have been developed to predict and correct for this bias^16–19^, including PRINT^20^, a recent deep learning model that substantially outperforms traditional k-mer-based approaches. However, like most existing methods, PRINT assumes a globally shared Tn5 bias profile across samples. Whether this assumption holds in bulk and single cell ATAC-seq experiments—especially across different tissues, protocols, or laboratories—remains unclear. If Tn5 cleavage bias varies across samples, global correction models may be insufficient, motivating the need for sample-specific approaches.

To investigate this question, we analyzed publicly available bulk and single-cell datasets spanning multiple tissues, technologies, and laboratories^3,21–25^. We found substantial variation in Tn5 cleavage bias across samples constituting a pervasive batch effect in both bulk and single-cell ATAC-seq data. These findings challenge the assumption of a global bias model and highlight the need for sample-specific correction.

These observations motivate us to develop TraceBIND (Transcriptional Regulation Analysis via corrected Epigenomic Bindings), a computational framework for high-resolution footprinting in bulk and single cell ATAC-seq data. TraceBIND leverages the mitochondrial-mapped reads from each sample to fine-tune cleavage bias predictions on a per-sample basis, generating nucleotide-resolution estimates of Tn5 activity. TraceBIND then applies a novel dynamic-window-dynamic-flank statistical scan to detect footprints with robust control of false positives, even under extreme variation in sequencing depths. By explicitly modeling sample-specific bias and integrating statistical evidence across multiple scales, TraceBIND improves the resolution and accuracy of regulatory element detection and provides a principled framework for refining peak calls into sub-peak regulatory features.

We performed simultaneous single-cell RNA and ATAC sequencing on aging rat kidney using the 10X Genomics single-cell multiome technology and analyzed the scATAC-seq portion with TraceBIND to investigate age-associated changes in chromatin accessibility and regulatory binding. Using TraceBIND, we observed a progressive increase in transcription factor footprinting with age, particularly at loci undergoing demethylation, even after controlling for overall accessibility changes. In contrast, nucleosome footprinting remained largely unchanged across age groups. To further enhance the interpretability of single-cell data, we integrated TraceBIND with existing tools such as Epitrace and chromVAR. By using sub-peak footprint features to inform cell age prediction and transcription factor activity inference, we demonstrate that TraceBIND can uncover subtle, cell-state-specific regulatory changes and provide new insight into the dynamics of transcriptional regulation during aging.

## Results

### Cross-sample variation in transposase cleavage bias introduces batch effects requiring sample-specific correction

Tn5 transposase cleavage bias is known to be sequence-dependent, but whether this bias varies across samples in large-scale bulk and single-cell ATAC-seq experiments remains unclear. To address this question, we analyzed 37 publicly available datasets spanning multiple technologies, tissues and laboratories^3,21-25^. Binding events in nuclear DNA can vary substantially across biological conditions, making it challenging to separate Tn5 transposase cleavage bias from biological variations in nuclear DNA. To overcome this, we leveraged insertion events in mitochondrial DNA (mtDNA), which are typically discarded in conventional ATAC analyses^19^. In the absence of histones, mtDNA better approximates deproteinized naked DNA^26^.

As in Hu et al.^20^, we computed a position-specific measure of Tn5 bias, defined as the fold enrichment of insertion count at each position relative to the average count across a local window centered at the position. Correlations of observed Tn5 bias in mtDNA between different cell types within the same sample is high, indicating that mitochondrial DNA accessibility does not vary across cell types and indeed serves as a stable in-sample control. Correlations of mtDNA-derived Tn5 bias is much lower across studies and laboratories, contrasting with the high correlations across cell types within the same sample (Fig. 1a-e and Supplementary Fig. 1, 2a-k). The same cross-sample variability was also observed when bias was quantified using k-mer preferences and raw insertion counts, confirming that Tn5 cleavage bias varies across samples, regardless of quantification method (Fig. 1f, g). These differences could not be attributed to variations in sequencing coverage, as downsampling to equal coverage across samples yielded similar results (Fig. 1h, i and Supplementary Fig. 3). The relationship between Tn5 bias and GC content varied across samples in a nonlinear and non-monotonic manner (Supplementary Fig. 2l). This cross-sample variability may be due to differences in experimental factors such as Tn5 concentration and the duration of exposure^27^.

**Fig 1.**
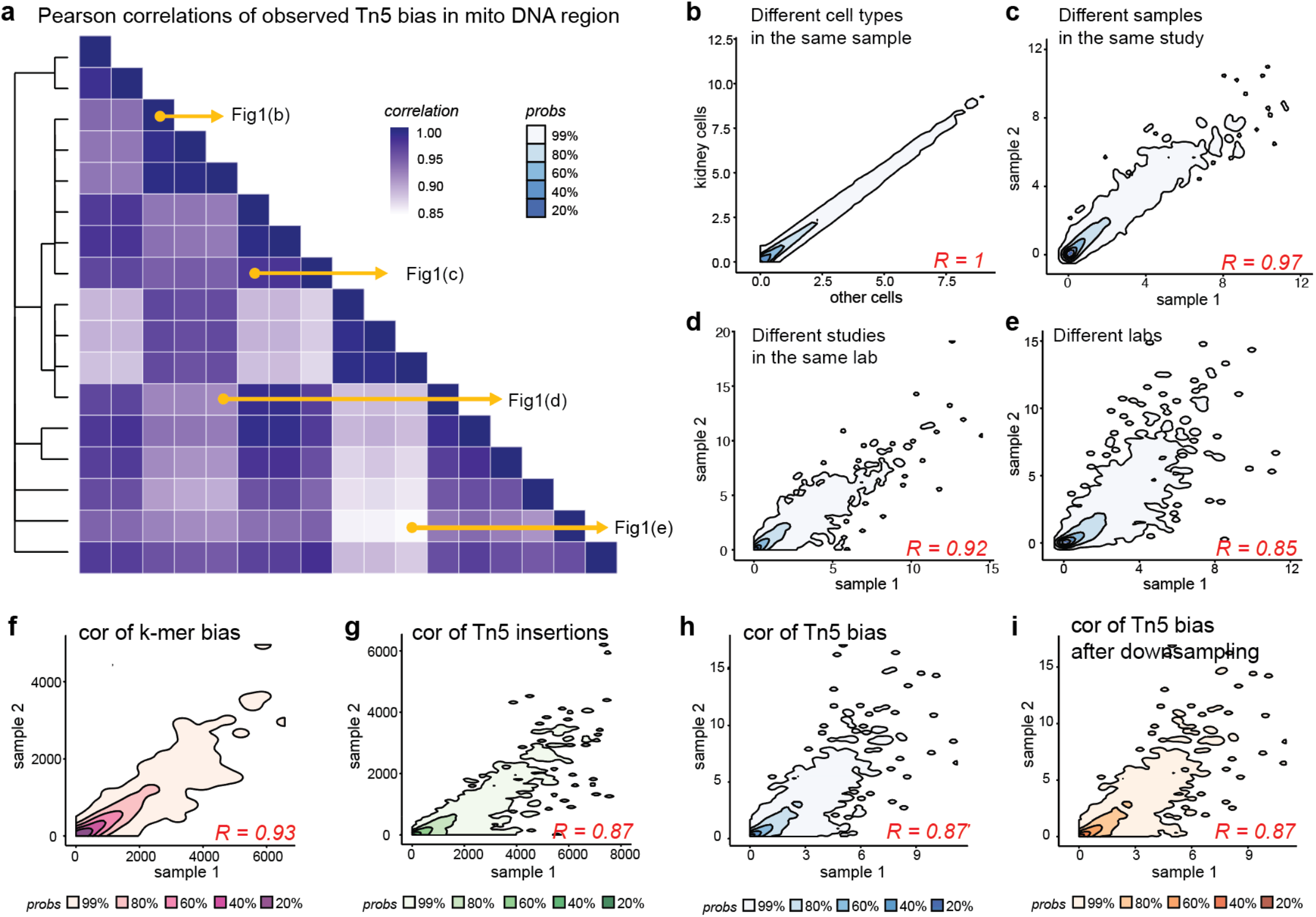
Sample-to-sample variations in Tn5 bias constitute severe batch effects. **a**, Correlations of observed Tn5 bias in mitochondrial DNA regions between samples reveal sample-to-sample variations, the dendrogram plot on the left shows relationships between samples at multiple levels: from the same study, different studies within the same lab, and between different labs. **b-e**, Density plots of Tn5 bias in mitochondrial DNA regions between 2 samples from different cell types, samples, studies and labs. **f**, Density plot of k-mer (k=5) bias between 2 samples in mitochondrial DNA regions. **g**, Density plot of observed Tn5 insertion counts between 2 samples in mitochondrial DNA regions. **h**, Density plot of observed Tn5 bias between 2 samples in mitochondrial DNA regions. **i**, Density plot of observed Tn5 bias between 2 samples in mitochondrial DNA regions after downsampled to the same coverage, indicating these differences cannot be explained by different coverage.

### TraceBIND: Robust multiscale footprint detection with sample-specific bias correction

Footprinting analysis in bulk and single cell ATAC-seq data requires accurate accounting for position-specific Tn5 transposase cleavage bias. A common approach is to estimate enzymatic cleavage bias from deproteinized naked DNA and apply these estimates to chromatin samples for correction^20,28,29^. However, our analysis has revealed substantial sample-specific variability in Tn5 bias, raising concerns about the effectiveness of global correction models. To address this, we developed TraceBIND *(Transcriptional Regulation Analysis via corrected Epigenomic binding)*, a two-step framework that first estimates sample-specific Tn5 cleavage bias, then identifies footprinting events through a multiscale adaptive scan. TraceBIND includes procedures for robust type 1 error control.

The Tn5 bias estimation step of TraceBIND builds on PRINT, a convolutional neural network for Tn5 bias prediction trained on deproteinized DNA^20^. We start with pretrained PRINT weights and perform sample-specific finetuning using the insertion counts from mtDNA (Fig. 2a). This allows us to leverage the expressiveness of convolutional neural networks in learning Tn5 bias at nucleotide resolution, while also capturing the sample-to-sample variations of this bias based on the limited complexity of the mitochondrial genome. Building on these predictions, TraceBIND computes a scan statistic to identify footprinting events using both observed insertions and bias-corrected expectations (Fig. 2b). For each nucleotide position and window size, the model quantifies both the p-value and effect magnitude for footprinting by comparing the window to a flanking region immediately bordering the window. For detecting weak footprint signals, larger flanking regions give a boost in power (Fig. 2c). However, an excessively large flanking window may overlap with neighboring footprints and confound signals (Fig. 2d). To balance sensitivity and specificity, TraceBIND utilizes a dynamic flank strategy, enabling sensitive identification of footprinting events in regions with complex chromatin landscape.

**Fig 2.**
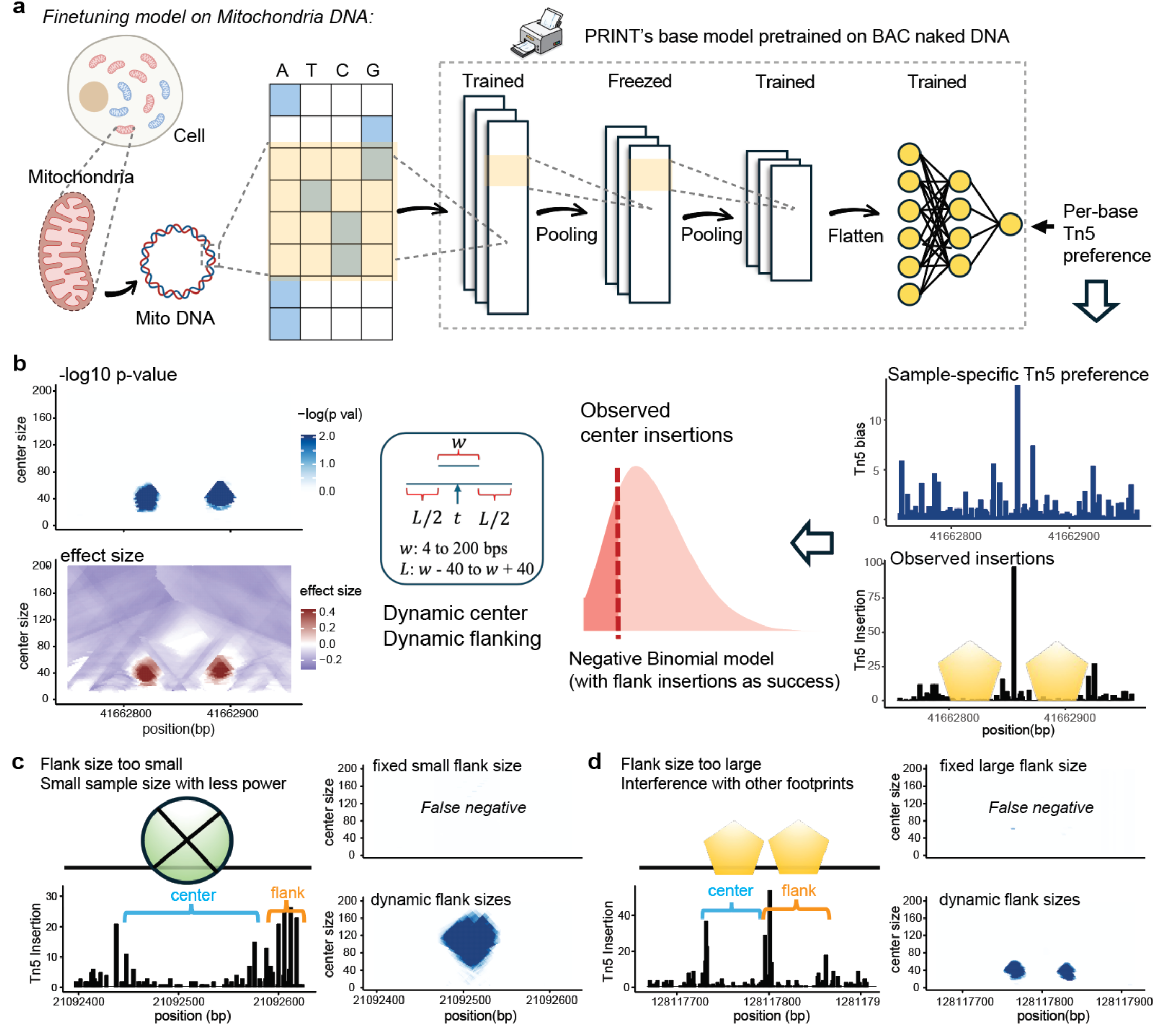
Overview of TraceBIND. **a**, Schematic Overview of TraceBIND workflow, (left to right), sample-specific Tn5 bias modelling, **b**, Schematic Overview of TraceBIND workflow (right to left), footprinting identification with dynamic flanking and center sizes. **c**, Illustration showing fixed small flank size will reduce power and increase false negatives due to insufficient background insertions. **d**, Illustration showing fixed large flank size will overlap with nearby footprints and increase false negatives.

TraceBIND systematically scans across center positions and window sizes, controlling for multiple testing, to identify non-overlapping binding events. The insertion counts from mtDNA further serves as an in-sample negative control for false positive thresholding. By repeatedly downsampling mtDNA reads to match the coverage observed in the search regions and applying our pipeline, we obtain sample-specific FDR-controlled p-value thresholds stratified by coverage, which is necessary because coverage can vary across orders of magnitude in ATAC-seq data.

### TraceBIND improves accuracy for footprint detection

We first evaluated methods on their specificity, i.e. their false positive rate in regions with no binding. We evaluated specificity using two types of negative control data: mitochondrial regions, which can be extracted from any ATAC-seq dataset, and naked DNA. Whereas mitochondrial regions extracted across ATAC-seq samples allow the assessment of sample-to-sample variation, ATAC-seq of naked DNA contains more sequence diversity and allows the assessment of whether TraceBIND’s strategy of finetuning on mitochondria reduces false positives on nuclear DNA. We will use mitochondria regions to evaluate bias prediction accuracy and naked DNA to assess the specificity of footprint detection^16,18,20^.

First, treating mitochondria as negative controls, we randomly partitioned the mitochondrial genome of each sample into training, validation, and test regions (Fig. 3a). Within the test regions, we downsampled to varying coverage to emulate the coverage levels typically observed for nuclear DNA and compared the observed Tn5 insertion rate with predictions from both PRINT and TraceBIND bias (Fig. 3b). Compared to PRINT, TraceBIND demonstrated a linear relationship and strong correlation with the observed Tn5 bias, which led to a significant reduction in false positives within the mitochondrial test regions across coverage levels (Fig. 3c and Supplementary Fig. 4a–f). This improvement is expected as TraceBIND builds on PRINT to finetune to sample-specific biases. We see that, across 37 samples, this sample-specific finetuning strategy improves Tn5 bias prediction. Notably, both PRINT and TraceBIND improve upon k-mer and position weight matrix (PWM) models, which are sample-specific but are not able to capture the complex sequence-specific biases (Fig. 3d and Supplementary Fig. 4g).

**Fig 3:**
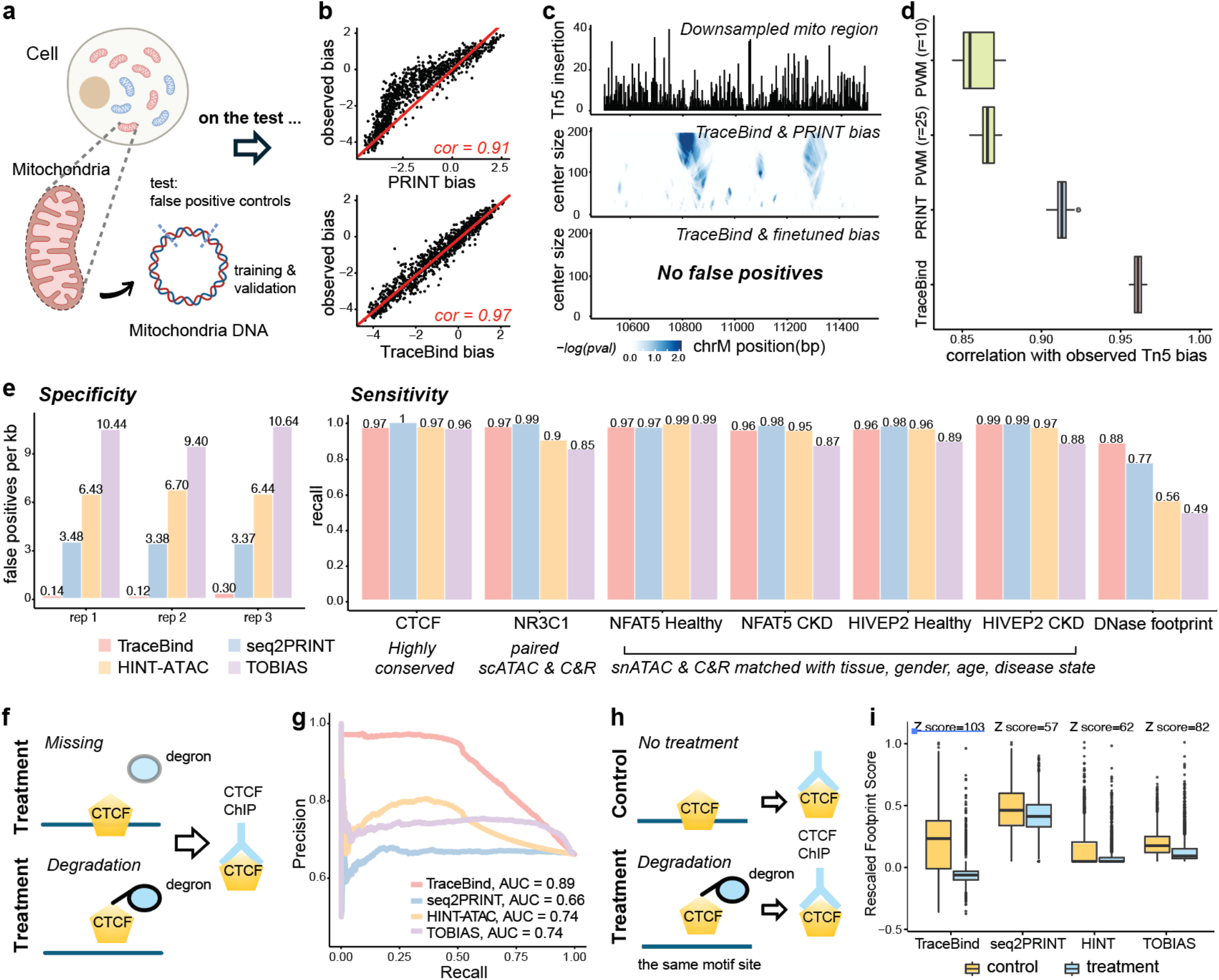
TraceBIND improves accuracy for footprint detection. **a**, Mitochondrial DNA regions were randomly split into train, validation and test sets; the benchmarking analyses were performed on the test data. **b**, Correlation between predicted and observed Tn5 bias on mitochondrial DNA test region for one sample. **c**, Footprint identification results on downsampled mitochondrial DNA test region, comparing Tn5 bias with and without TraceBIND finetuning. **d**, Boxplot of correlations between predicted and observed Tn5 bias on mitochondrial DNA test region across 37 samples. **e**, Benchmarking of footprint identification methods on naked DNA and scATAC-seq data with matched ChIP-seq/CUT&RUN as ground truth. **f**, Schematic Overview of CTCF-Degron experiments workflow. **g**, Benchmarking of specificity and sensitivity on CTCF-Degron experiments. **h**, Schematic Overview of comparisons between control and CTCF-Degron treatment at degron-depleted motif sites. **i**, Footprint prediction scores at degron-depleted motif sites in control and treatment. Z scores from Wilcoxon signed-rank tests were computed, where larger Z scores mean larger differences between control and treatment.

Next, we benchmarked specificity on public ATAC-seq data from naked DNA^30^. We applied TraceBIND to finetune PRINT cleavage bias estimates using the insertion profile from the mitochondria of this sample, then detected footprints on the nuclear genome. We see that TraceBIND substantially improved on existing methods^16,18,20^, reducing false positive rate by orders of magnitude (Fig. 3e left and Supplementary Fig. 4i) across the 3 samples: Compared to TOBIAS, on average, the false positive rate was reduced by > 60 fold. Compared to HINT-ATAC, the false positive rate was reduced by > 30 fold. Compared to PRINT, the false positive rate was reduced by > 20 fold. This proves that our strategy of using the mitochondria data of each sample for finetuning indeed substantially reduces the false positive rate on the nuclear genome.

Next, we considered sensitivity. To evaluate sensitivity, we harnessed publicly available ChIP-seq, CUT&RUN (C&R), and DNase-seq data, and evaluated recall of the binding sites detected by these technologies. Since chromatin state and TF binding are highly context specific, we restricted our analysis to cases where the validation data are generated under conditions that closely reflect the biology of the ATAC-seq experiments^31^ on which footprinting was performed.

Thus, we focused on highly conserved TFs^3^ or selected datasets in which the ChIP-seq/CUT&RUN and scATAC-seq experiments are matched for tissue, gender, age, and disease state^22,25,32^. We identified 7 such ChIP-seq/CUT&RUN ground truth data sets, and for each, computed the recall of their identified binding events in the matched pseudobulked scATAC-seq data. The results show that TraceBIND achieved a high recall of experimentally supported TF binding events, with sensitivity that was on par or better than existing methods despite the orders of magnitude decrease in false positive rate (Fig. 3e right and Supplementary Fig. 4j). DNase-seq is widely regarded as the gold standard technology for TF footprinting^33,34^, with previous studies showing concordance between binding sites identified by ATAC-seq and DNase-seq^16,35^. Using DNase-seq-derived binding sites from kidney proximal tubule (PT) cells as ground truth^28^, we evaluated the recall of footprinting predictions from pseudobulked scATAC-seq of PT cells. The result showed that TraceBIND recovered the highest number of DNase footprints, 10% higher than the next best method and significantly improving upon existing state-of-the-art methods (Fig. 3e right and Supplementary Fig. 4h).

To further assess the balance between sensitivity and specificity, we evaluated the performance of each method in scATAC-seq data from HCT116 cells before and after degron-induced CTCF depletion, with paired CTCF ChIP-seq for each condition serving as ground truth^3^. In degron-treated condition, a total of 4,684 CTCF motif sites that overlapped with summits of both control and treatment ChIP-seq were labeled as true positives, while 2,393 motif sites that bound in the control ChIP-seq but not in treatment ChIP-seq were considered as degron-depleted and labeled as true negatives (Fig. 3f). TraceBIND achieved a significantly higher area under the precision-recall curve (AUPR) than existing methods (Fig. 3g), suggesting it could predict condition-specific bindings. Moreover, at degron-depleted CTCF motif sites (Fig. 3h), TraceBIND showed the most significant differences in prediction scores between control and treatment (Fig. 3i and right and Supplementary Fig. 5), indicating its sensitivity to biologically meaningful changes in binding.

### TraceBIND captures gradual loss in footprint strength across increasing levels of nucleosome depletion

We next investigated whether TraceBIND’s p-values and estimated effect sizes can be used as quantitative measures of footprinting strength. To this end, we performed DEFND-seq, which is a modification of single-cell multiome (RNA + ATAC) sequencing that employs lithium diiodosalicylate to deplete nucleosomes and expose DNA to Tn5 transposase^36^. We hypothesized that increasing concentrations of lithium would deplete more nucleosomes and serve as a marker of nucleosome stability. We analyzed pseudobulked scATAC-seq from our DEFND-seq libraries using three conditions: chromatin in its original state without lithium treatment, chromatin after weak lithium treatment, and chromatin after strong lithium treatment. To ensure comparability, we limited our analysis to peaks that are shared across the three samples from the same age group and downsampled cell numbers for high coverage conditions to match samples in total coverage. Footprint counts were highest in the no-depletion dataset, followed by the weak-depletion and then the strong-depletion datasets, showing a clear monotone decrease in footprint number with increasing depletion levels across footprint widths (Fig. 4a). Similar trends were observed in the effect sizes of the detected footprints: Effect sizes for footprints wider than 120 bp, which contain putative nucleosome binding sites, were highest in the no-depletion dataset, followed by weak-then strong-depletion DEFND-seq datasets, with significant differences between all conditions by Wilcoxon signed rank test (Fig. 4b). Fig. 4c shows a representative region, where both the significance (left) and effect size (right) of footprinting signals decrease with increasing levels of nucleosome depletion. This serves as proof of principle that TraceBIND effect sizes can serve as a quantitative proxy for footprint strength.

**Fig 4:**
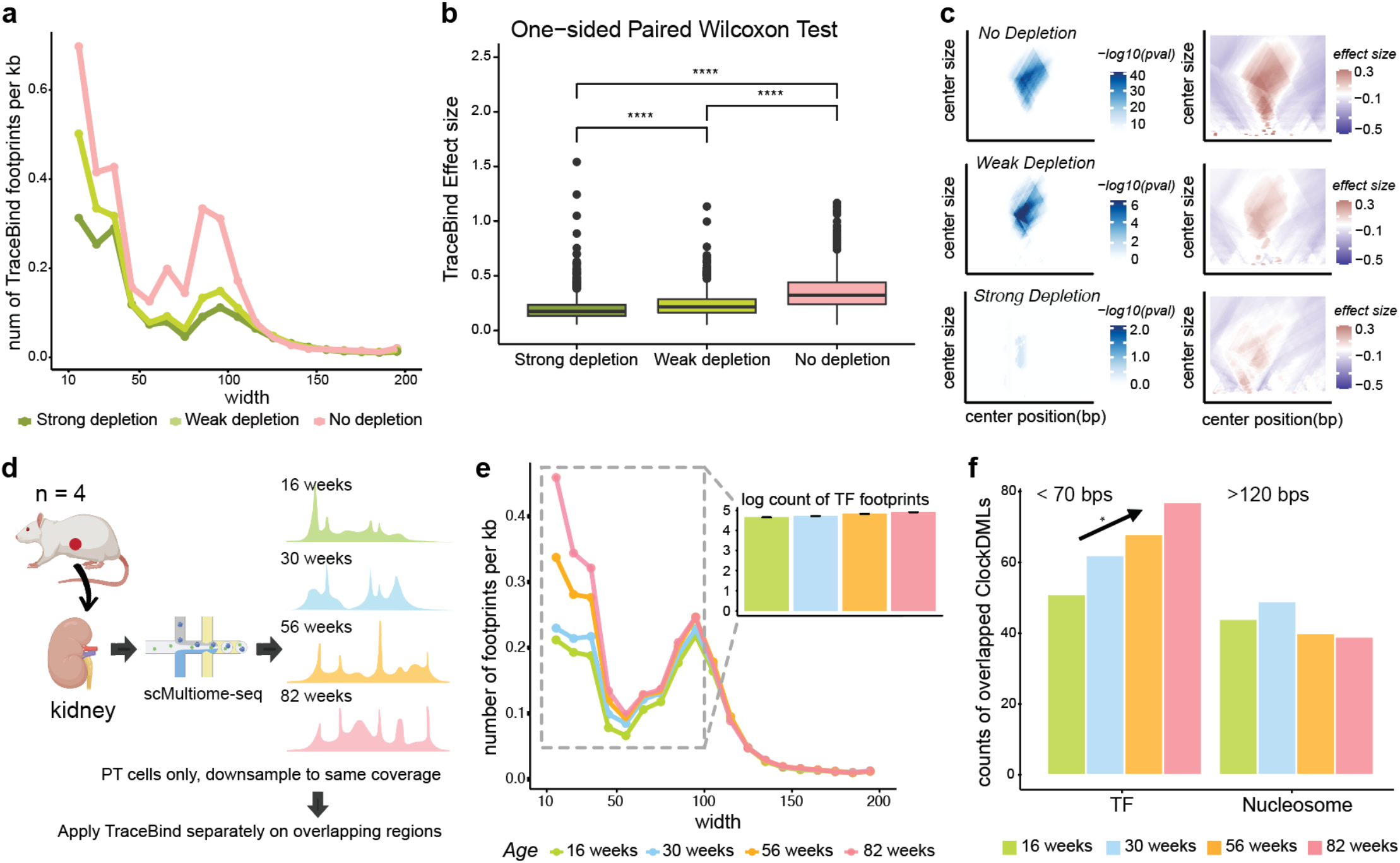
TraceBIND gives quantitative measures of footprint strength and uncovers age-related footprinting signatures. **a**, Counts of footprints for different widths across different depletion levels of nucleosomes. **b**, Predicted effect sizes of nucleosomes across different depletion levels of nucleosomes. **c**, A nucleosome depletion example showing signals across different depletion levels of nucleosomes. **d**, Schematic overview of aging experiments and TraceBIND analyses. **e**, Counts of footprints for different widths across four age groups of rat kidney proximal tubule cells. **f**, Counts of TF and nucleosome footprints overlapping with age-associated differentially methylated loci.

### TraceBIND uncovers age-related footprinting signatures and refines cell age prediction in kidney

Across tissues and species, age-associated chromatin changes have been reported^37–40^. Leveraging the improved footprint accuracy enabled by TraceBIND, we investigated age-related changes in chromatin accessibility and protein binding from single-cell multiome (RNA + ATAC) sequencing of aged rat kidney, sampled at four time points across the rat lifespan: 16, 30, 56, and 82 weeks (Fig. 4d and Supplementary Fig. 6a, b). We focused on proximal tubule cells, a key cell type implicated in chronic kidney disease and known to initiate injury and inflammation responses. Our data set includes 9,278 proximal tubule cells across the four age groups, each sequenced to a median of 11,209 reads. A total of 146,971 peaks were detected across the four age groups. To ensure comparable coverage across samples, we randomly downsampled cell numbers for each age group.

We observed a monotonic increase with age in both the percentage of the genome in accessible peaks and in the average peak count per cell (p-value of pearson correlation with age = 0.03, 0.04 respectively, Supplementary Fig. 6c, d). This trend aligns with previous studies reporting that aging is associated with loss of cell identity and a depletion of repressive epigenetic marks^38,40^, resulting in an increased proportion of accessible chromatin^39,41^. We next asked whether transcription factor (TF) binding and nucleosome occupancy exhibit per-base changes with age, after adjusting for the overall increases in peak count. To isolate this effect, we restricted analysis to peaks shared across all age groups and randomly downsampled cell numbers to equalize aggregate coverage between samples (Fig. 4d). TraceBIND was then applied with FDR controlled at 5%.

There are two modes in the width distribution of the detected footprints: at ∼10 and ∼100 base pairs. Given this, we attribute the footprints with width >= 120 base pairs as “putative nucleosome footprints”^42^, and the footprints <= 70 base pairs as “putative TF footprints”. Footprints that >= 70 bps and <= 120 bps are interpreted as a mixture of submononucleosome^43^, structured DNA elements^44^ and multi-TF complexes^45^. Across these shared peak regions, we observed significant, monotonic increase in the frequency of TF footprinting with age (p-value of permutation = 0.04). Because coverage was equalized, this trend reflects a true increase in binding frequency, rather than sequencing depth or total availability of accessible chromatin. In contrast, the frequency of nucleosome footprints remained relatively stable across ages, where the increase is insignificant (Fig. 4e).

DNA methylation is known to change as cells age and has been used to construct accurate molecular clocks of aging^37–40^. We hypothesize that the progressive increase in TF footprint frequency may be due to age-associated loss of repressive marks, such as DNA methylation. To test this, we examined TraceBIND footprints overlapping with age-associated differentially methylated loci (DMLs)^46^, which are DNA loci that are known to be methylated in early life and progressively de-methylated with age in general^38,47^. We found that the proportion of age-associated DMLs residing in putative TF footprints increases significantly and monotonically with age (p-value of permutation = 0.04). In contrast, the proportion of age-associated differentially methylated loci residing in putative nucleosome footprints showed no consistent trend (Fig. 4f), This is consistent with the notion that DNA methylation disrupts TF-DNA binding but does not directly disrupt nucleosome positioning. This trend is also observed in an independent age-associated DMLs dataset that exclusively contains hypomethylation sites^48^ (Supplementary. Fig 6e).

### TraceBind refines TF regulatory analysis and cell age prediction in kidney scATAC-seq

We then sought to investigate the TF activity changes in kidney proximal tubule cells from a young adult rat (30 weeks) to a late-stage aged rat (82 weeks). ChromVAR^49^, a widely used tool for inferring TF activity from scATAC-seq data, quantifies variability in chromatin accessibility at motif sites for each single cell to estimate cell-specific activity of TFs in an unbiased manner. However, an ATAC peak often contains multiple TF motif sites, especially those of closely related TFs, and standard analyses cannot distinguish between bound and unbound sites, potentially confounding TF activity inference and leading to spurious predictions. To improve specificity, we restricted chromVAR motif input to only those motif sites that overlapped with TF footprints identified in TraceBIND analysis of cell type-level pseudo-bulk data. By integrating footprint information into chromVAR analysis, we observed an age-associated upregulation of AP-1 family members (Fig. 5a), consistent with prior reports of increased AP-1 activity during aging in various tissues, including kidney^50–53^. In contrast, the standard chromVAR analysis on all motif sites suggested a decline in AP-1 activity with age which does not align with established findings. This analysis demonstrates that focusing on actively bound motif sites detected by TraceBIND could enhance the biological relevance and accuracy of TF activity scores. Similar trends were observed across different age strata (Supplementary Fig. 6f-h).

**Fig 5:**
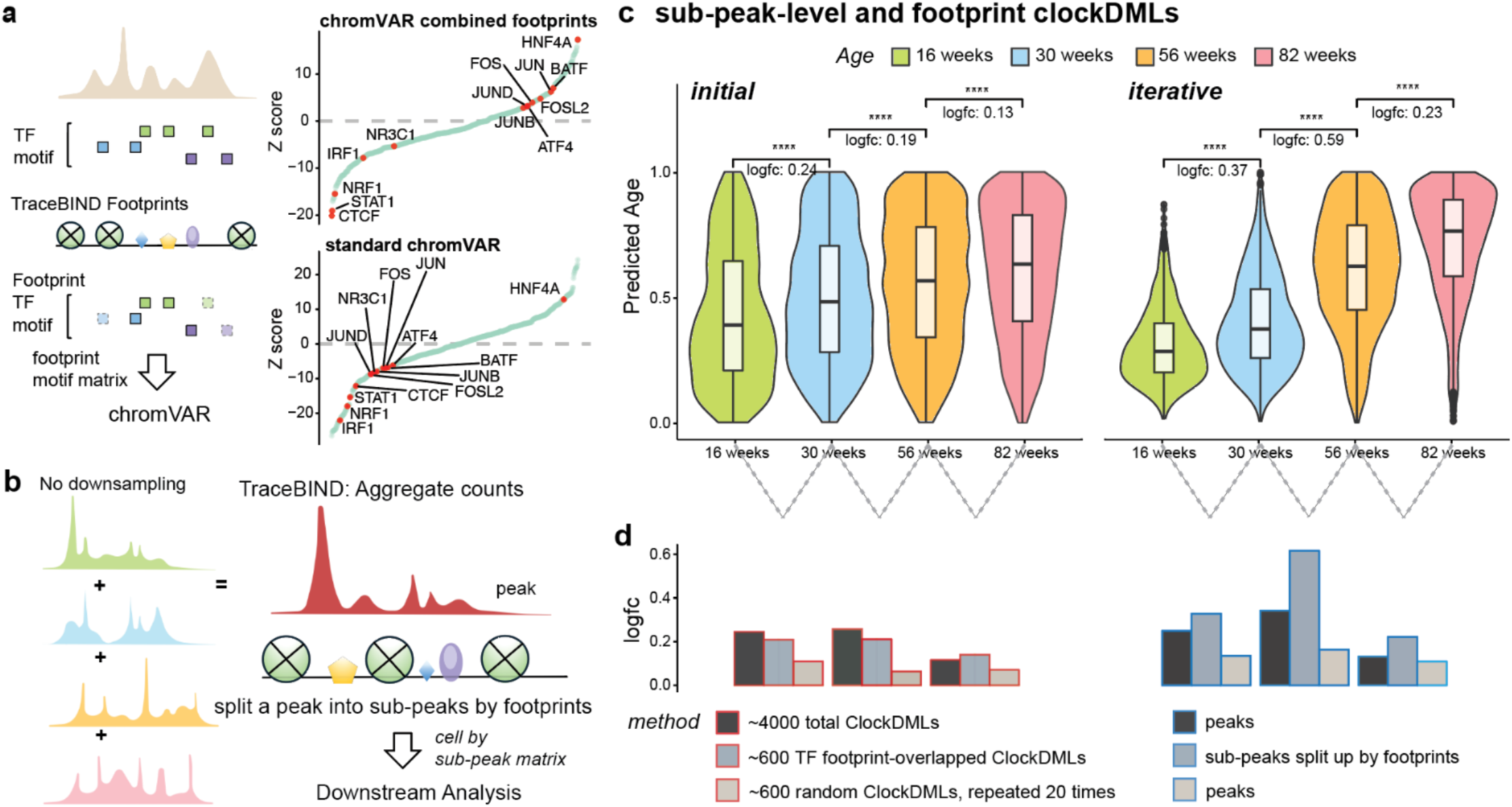
TraceBind footprinting improves TF activity analysis and cell age predictions in kidney. **a**, Schematic overview, and results of footprint-refined chromVAR, comparing standard and footprint-refined TF activity inference. Positive Z-scores indicate higher TF activity in the aged group. **b**, Schematic overview of sub-peak-level analyses. **c**, Age prediction using footprint-overlapping age-associated differentially methylated loci at sub-peak-level. **d**, Comparison of distinctions across ages between standard peak-level, footprint-informed sub-peak-level, and random DML subsets peak-level.

Age-associated DMLs were recently harnessed to estimate the mitotic age of cells from scATAC-seq data^46^. Epitrace^46^ reduces the data to a peak-by-cell matrix, counting the peaks that overlap with age-associated DMLs to form an initial estimate of cell age, then refining this estimate through iteratively including newly identified age-associated peaks. Motivated by our finding of age-associated increase in TF footprinting frequency, we sought to enhance Epitrace by first using TraceBIND to detect footprints across all 146,971 peaks, then partitioning each peak into finer sub-peak regions (Fig. 5b). This generates higher-resolution features for cell age estimation (Supplementary Fig. 7a, b).

We applied this strategy to the scATAC-seq data of kidney proximal tubule cells from aging rats, where we expect to observe a monotone upward shift in the distribution of estimated cell ages from 16 to 82 weeks. First, we compared the initial cell age estimates derived from simply counting overlaps with ClockDMLs, prior to the iterative update step. These initial cell age estimates indeed increased monotonically with age (Fig. 5c left). Notably, restricting to the 639 ClockDMLs residing within TraceBIND TF footprints out of the 4,170 total

ClockDMLs within peaks recovered almost the same age-related shift (Fig. 5d left, and Supplementary Fig. 5f). This indicates that footprint-based features can capture aging signals with far fewer loci. In contrast, randomly selected sets of 639 loci yielded unstable, less distinct, and sometimes incorrect age predictions (Fig. 5d left and Supplementary Fig. 7c-d).

Next, we performed the Epitrace iteration step using both peak-level features and TraceBIND-derived sub-peak-level features. Fine-tuning based on sub-peak-level features produced a much stronger monotonic trend with aging compared to fine-tuning based on peak-level features alone (Fig. 5c and 5d right). This suggests that TraceBIND-derived sub-peak features preserve more nuanced regulatory signals, enabling more accurate cell age estimation.

## Discussion

Footprint analysis on ATAC-seq data holds the promise to reveal direct protein–DNA interactions. By detecting localized patterns of protection from transposase activity, footprinting enables the identification of transcription factor binding sites and nucleosome positioning at base-pair resolution, providing fine-scale mechanistic insights into how regulatory elements control gene expression. However, its application and wide-spread adoption has been hindered by technical noise, especially those due to Tn5 cleavage bias. We show here that Tn5 cleavage bias varies substantially between samples, introducing a previously underappreciated batch effect and challenging the assumptions of current footprint methods that rely on Tn5 bias estimated from deproteinized naked DNA controls.

We developed TraceBIND, a statistical model and analysis framework that accurately estimates Tn5 bias in a sample-specific manner from ATAC-seq data and enables more precise identification of transcription factors and nucleosome footprints. A key innovation of our approach is the leveraging of mitochondrial DNA as an in-sample negative control for Tn5 bias prediction. By harnessing the recent PRINT bias model and incorporating mitochondrial reads into the bias correction step, our framework captures sample-specific cleavage patterns without the need for external controls, improving the robustness of bias correction across tissues, protocols, and sequencing depths. We use a diverse set of gold-standard validations to show that TraceBIND maintains high sensitivity while markedly reducing the false positive rate. Remarkably, while TraceBIND was not trained on labeled TF binding data, it outperforms existing supervised tools.

We applied TraceBIND to a single cell study of aging, which is known to be associated with widespread epigenomic change. While much has been reported about age-associated shifts in DNA methylation and global chromatin accessibility, transcription factor footprinting remains underexplored due to technical limitations. Using kidney proximal tubule cells from rats across 4 age groups spanning the lifespan, we observed a monotonic increase in TF footprinting with age, even after accounting for changes in global accessibility. In contrast, nucleosome footprinting is relatively stable with age. As expected, the increase in TF footprinting is especially evident at loci undergoing hypomethylation, validating our approach and connecting age-associated demethylation with age-associated increase in TF binding.

Although ATAC-seq footprinting offers high-resolution insight into regulatory activity, it is inherently coverage-intensive and challenging to apply directly to sparse single-cell data. In scATAC-seq studies, pseudobulking across predefined cell states is typically required, limiting footprinting analyses to well-sampled populations. However, with the increasing scale of scATAC-seq studies now spanning tens to hundreds of samples, cell numbers are becoming sufficient to support footprinting at fine cell state resolution. TraceBIND integrates with existing single-cell analysis pipelines such as chromVAR and Epitrace, improving TF activity inference and cell age estimation by incorporating precise, footprint-informed features. Overall, TraceBIND enables unbiased, high-resolution mining of these expanding datasets to uncover cell type– and state–specific regulatory signatures.

## Methods

### Aged rat kidney cortex samples

Male Wistar rats aged 14 to 80 weeks were purchased from Janvier laboratories and housed at the University of Pennsylvania institutional animal care facilities for two weeks prior to euthanasia. Rats were euthanized with carbon dioxide and kidneys were removed. Kidneys were dissected to separate the cortex and medulla and then snap-frozen in liquid nitrogen for future single-cell studies.

### scMultiome-seq and DEFND-seq from rat kidney cortex

Nuclei were prepared following an adapted protocol based on procedures from Olsen et al.^36^. In brief, 25 mg snap-frozen rat kidney cortex was placed in 2 ml of tissue lysis buffer (10 mM Tris-HCl pH 7.4, 10 mM NaCl, 3 mM MgCl_2_, 0.1% IGEPAL CA-630 (Milipore Sigma, 18896), 0.005% digitonin (Invitrogen, BN2006), 1 mM DTT, 0.5 U/μL Protector RNase Inhibitor (Milipore Sigma, 3335402001)). The tissue was then dissociated with a GentleMACS Dissociator (Miltenyi Biotec) using the Multi_E_01 and Multi_E_02 programs. After dissociation, 1 mL PBS Buffer (PBS with 1% BSA (MACS® BSA Stock Solution, Miltenyi Biotec, 130-091-376), 1 mM DTT, 0.5 U/μL Protector RNase inhibitor) was added to quench the lysis reaction. The lysate was strained through a 40 μm filter and spun down at 400*g* for 4 min at 4 °C. Nuclei were resuspended in 1 ml PBS buffer and spun again at 400*g* for 4 min at 4 °C to remove residual lysis buffer. Isolate nuclei were used to generate either scMultiome or DEFND-seq libraries as follows:

1. For scMultiome samples, nuclei were resuspended in 1 ml PBS buffer and spun again at 400*g* for 4 min at 4 °C. Finally, nuclei were resuspended in 150 μL 10x Genomics Diluted Nuclei Buffer (1X Nuclei Buffer (10x PN 2000207), 1 mM DTT and 1U/μL Protector RNase Inhibitor).
2. For DEFND-seq samples, nuclei were resuspended in 200 μL PBS buffer supplemented with lithium 3,5-diiodosalicylate (Li; Millipore Sigma, D3635-5G, dissolved in nuclease-free water) at a concentration of either 12.5 mM Li (strong depletion) or 6 mM Li (weak depletion) and incubated on ice for 5 min to deplete nucleosomes. Immediately after the incubation, 800 μL PBS buffer was added to quench the reaction. The nuclei were then centrifuged at 400*g* for 4 min at 4 °C and resuspended in 500 μL wash buffer (10 mM Tris-HCl pH 7.4, 10 mM NaCl, 3 mM MgCl_2_, 1% BSA, 0.1% Tween-20, 1 mM DTT, 0.5 U/μL Protector RNase inhibitor) and spun again at 400*g* for 4 min at 4 °C. Finally, nuclei were resuspended in 150 μL 10x Genomics Diluted Nuclei Buffer (1X Nuclei Buffer (10x PN 2000207), 1 mM DTT and 1U/μL Protector RNase Inhibitor).

Nuclei were then used to generate single-cell libraries according to Chromium Next GEM Single Cell Multiome ATAC + Gene Expression user guide (protocol CG000338 Rev F) with corresponding reagents (10x Genomics PN-1000282, 1000280, 1000190, 1000234, 1000212 and 1000215).

ATAC and GEX libraries were sequenced on Illumina NovaSeq X platform with 151 cycles for read 1, 10 cycles for index 1, 24 cycles for index 2, and 151 cycles for read 2. All libraries were sequenced with 1% PhiX. **scMultiome and DEFND-seq data processing:** Raw BCL sequence files were demultiplexed to generate Fastq files using BCL Convert Software (v4.3.13) according to suggestions from 10x Genomics^54^. For ATAC libraries, read 1 was trimmed to 8 cycles; for GEX libraries, read 2 was trimmed to 10 cycles. The ATAC reads were processed with Cell Ranger ATAC (v 2.1.0) using the GRCr8 reference genome^55^. As GRCr8 does not have a mitochondrial reference, we used the mitochondrial genome from mRatBN7.2. RNA reads were processed with Cell Ranger (v 8.0.0) to generate gene count matrices based on the same reference genome. RNA count matrices were aggregated with Cell Ranger (with normalize = “none”) and imported into R to create Seurat objects (v 5.1.0). Cells were filtered based on the following quality control criteria: between 400 and 7000 RNA features and less than 30% mitochondrial counts. Doublets were removed by scDblFinder^56^ with removeUnidentifiable = TRUE. The gene expression matrix was then normalized using SCTransform. Harmony^57^ was used to integrate expression data based on the SCTransform Pearson residual matrix. 50 dimensions from Harmony reduction were used for FindNeighbors and resolution = 0.6 was used to perform FindClusters functions in Seurat^58^. Cell type annotations for scMultiome-seq and DEFND-seq were based on expression of lineage-specific genes.

### Computation of p-values and effect sizes

Let *w* be footprint center region size, *L* be flanking region size (in base pair). Let 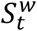 be counts of Tn5 insertions within the window of size *w* centered at *t*, 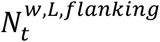 be counts of Tn5 insertions within (*t* - *w*/2 - L, *t* - *w*/2 - 1) or (*t* + *w*/2 + 1, *t* + *w*/2 + L). Suppose that when there is no footprinting, using flanking insertions as baselines:

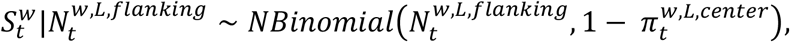

Where

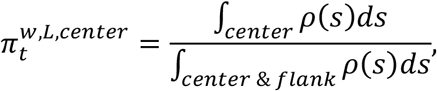

and *ρ*(*s*) is the predicted Tn5 insertion preference at base *s*. We test the hypotheses:

*H*_0_: there is no footprinting with width *w* centered at position *t*,

*H*_1_: there is such footprinting with width *w* centered at position *t*. where the p-value for each test is:

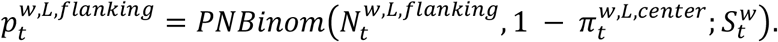

Following PRINT^20^, we perform two separate tests: one comparing the center to the left flank and one comparing the center to the right flank. To reduce false positives from asymmetric insertion patterns, we retain the larger of the two p-values.

We apply the Cauchy Combination Method to combine p-values 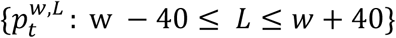 to obtain 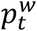, where weights are decided based on an exponential decay function exp(−|*L* − *w*|). Then, to identify putative footprints, we apply multiple test adjustments to 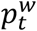 for each *t, w* within peak region (Benjamini-Hochberg). If the null and alternative hypotheses test for the presence of any footprint within a peak region, Benjamini-Yekutieli (BY) correction is applied for multiple testing adjustment.

### Identification of footprints

We identify footprints within peak regions by selecting non-overlapping regions with the smallest multiple-test-adjusted p-values, representing the most statistically significant footprinting signals. For each significant region, the width and position are further determined based on the location of maximum effect size within that region.

### Getting Tn5 insertion counts

Following procedures by PRINT, fragment ends were offset by +4/−4 bp to align with the center of the 9 bp staggered cut generated by Tn5 transposition.

### Derivations of mitochondria Tn5 fragments

Mitochondria reads were extracted from bam files by samtools^59^ and sinto^60^ with the following parameters: samtools view -b, samtools sort -o, samtools index and sinto fragments -b -f --collapse_within.

### Estimations of sample-to-sample variations in Tn5 observed bias in mitochondria regions

After obtaining mitochondria fragments and Tn5 insertions for each position along mitochondrial genome, we calculated observed Tn5 bias for each base pair *t* as:

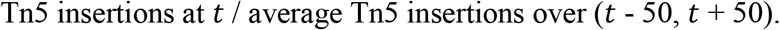

To assess variation between samples, we computed the Pearson correlation of observed Tn5 bias profiles in mitochondrial regions between all pairs of samples.

### Estimation of dispersion in negative binomial models across different cell types within the same group and across different groups

For Tn5 insertions *X*_*i*_ at base pair *i* in mitochondria region, suppose that *X*_*i*_ ∼ *NBinomial*(*N*_*i*_*ρ*_*i*_, *ϕ*), where *N*_*i*_is the average number of insertions across a local window around position *i*, and *ρ*_*i*_represents the observed bias at position *i* estimated using all cell types. The term *N*_*i*_*ρ*_*i*_ could thus be viewed as the mean of the negative binomial distribution, while *ϕ* denotes the dispersion parameter. To evaluate whether the sample-level observed bias (*ρ*_*i*_) adequately captures cell type-specific bias, we fit the model 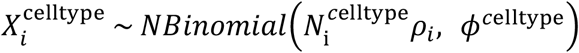 for each cell type within each sample. If sample level bias *ρ*_*i*_ could represent each individual cell type, the estimated dispersion *ϕ*^*c*elltype^ should be small. We also fit the model 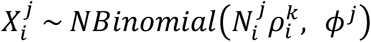 for each sample *j*, using 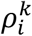 estimated in the other sample *j* (*j* ≠ *k*). If the substantial sample-to-sample variations in Tn5 bias exist, the estimated dispersion *ϕ*^*j*^ should be large.

### Estimations of sample-to-sample variations in Tn5 observed bias in mitochondria regions after downsampling

To control for coverage differences when comparing observed Tn5 bias between samples, we perform Binomial sampling to downsample Tn5 insertions at each position. Specifically, for the sample with higher coverage, insertions were downsampled using binomial sampling Binomial(*n, p*), where *n* is the observed number of Tn5 insertions at a given position and *p* is the ratio of average coverage between the two samples. After downsampling, we recalculated the observed Tn5 bias and computed the Pearson correlation between the two samples. This procedure was repeated 50 times for each sample.

### Tn5 bias Model structure

The model architecture follows the PRINT framework and consists of three convolutional-max pooling blocks followed by two fully connected layers. Each block uses 32 filters of width 5, with ReLU activation, same padding, and a stride of 1. The dense layers output 32 and 1 units, with ReLU and linear activations, respectively. The input is a one-hot encoded ±50 bp DNA sequence centered on a given mitochondrial position, and the output is the log transformed, rescaled observed Tn5 bias. Mitochondrial regions were randomly split into training (85%), validation (10%), and test (5%) sets.

### Fine-tuning PRINT with mitochondrial Tn5 reads

The model was finetuned on the training set with the second convolutional layer frozen. This freezing strategy preserves mid-level sequence features learned from the original PRINT model, which could capture generalizable patterns of Tn5 cleavage, while allowing early and late layers to adapt to sample-specific signal and output scaling. Hyperparameters were optimized using the validation set, and final performance was assessed on the test set. Training used mean squared error as the loss function, the Adam optimizer with default settings, a batch size of 64, and early stopping based on validation loss. The model was implemented in Keras.

### Benchmarking with other Tn5 bias models

The benchmarking was performed on the test set using several alternative bias methods include PRINT, k-mer (k = 3, 5 and 7), and PWM (radius = 10, 25). For k-mer models, foreground and background frequencies of all possible k-mers were computed from the mitochondrial training data. For PWM models, we calculated foreground and background base frequencies within a ±10 or ±25 bp window around each insertion site in the training set and used these to construct a position weight matrix (PWM) representing Tn5 insertion bias for the test set. For the PRINT neural network model, Tn5 bias in mitochondrial test regions was predicted directly from the DNA sequence input. Both predicted and observed Tn5 bias values were log-transformed prior to Pearson correlation calculation, as the data were heavily left skewed.

### Stratified false positive thresholding by mitochondria

Because sequencing coverage varies widely across the genome, we establish a data-driven threshold for significance. The mitochondrial test region is used as an in-sample negative control. To account for differences in sequencing depth, mtDNA reads are repeatedly downsampled to match the average coverage levels observed in the genomic search regions. Specifically, Tn5 insertions are downsampled using binomial sampling, Binomial(*n, p*), where *n* is the observed number of insertions at a given position in the mitochondrial test region, and *p* is the ratio of the target coverage level to the average mitochondrial coverage. Our pipeline is then applied to these downsampled sets to empirically estimate p-value thresholds corresponding to a target FDR, stratified by average coverage. The final threshold is the minimum of the model-based and stratified FDR-adjusted p-values.

### Validations and benchmarking with existing methods

The seq2PRINT model is designed to compute TF and nucleosome binding scores for every 10-base-pair bin within peaks. For each bin, we used the maximum of TF and nucleosome binding scores as the final score. Bins with scores ≥ 0.5 were considered as binding sites. Consecutive positive bins were merged and counted as a single binding event. For TraceBIND, the binding\ events are identified at a false discovery rate of 10%. HINT-ATAC^16^ and TOBIAS^18^ were applied using their default parameters.

### Naked DNA

To evaluate the specificity of each model, we also tested all four models on the naked DNA data^30^, where no true binding events are expected. The number of predicted binding events are used to represent the false positive predictions.

### ChIP/CUT&RUN

For sensitivity evaluation of each model, we used ChIP-seq/CUT&RUN^3,22,25,32^ as ground truth to validate predicted binding events. Given the dynamic nature of TF binding across cell states, we carefully selected scATAC-seq datasets that were well-matched to the corresponding ground truth data. NR3C1 CUT&RUN and scATAC-seq were performed on the same sample, while CUT&RUN and scATAC-seq for NFAT5 and HIVEP2 were obtained from the same study and matched with gender, age and disease state. CTCF ChIP-seq data on kidney, representing a highly conserved TF, were obtained from ENCODE. Peaks with low average coverage, defined as fewer than one insertion per base pair, were excluded from the analysis. Only TF binding sites overlapping with scATAC-seq peaks were retained for evaluations.

### DNase Footprints

We took the intersection of DNase footprint sites^28^ identified from three healthy proximal tubular samples (adjusted p.value <= 1e-4) as ground truth, which were downloaded from https://www.vierstra.org/resources/dgf. In scATAC-seq data, only proximal tubular cells were included as input for footprinting detection. Similarly, peaks with low average coverage (< 1 insertions per bp) were excluded from the analysis.

### CTCF Degron

Paired CTCF ChIP-seq and scATAC-seq datasets^3^, performed before and after CTCF degron treatment, were downloaded from ENCODE. Given that CTCF motif sites are well characterized, we retained only those ChIP-seq binding sites that overlapped known CTCF motif sites. Motif sites with consistent binding by ChIP-seq in both conditions were labeled as positives, while those with binding lost after treatment by ChIP-seq were labeled as negatives. To investigate changes before and after degron treatment, we included only scATAC-seq peaks shared between control and treatment with mean insertions per base pair > 0.2. We then evaluated each method’s predictive performance by calculating the area under the precision-recall curve (AUPR) based on predicted scores at these TF binding sites. For CTCF-depleted sites, we assessed each method’s sensitivity to binding loss by computing Z-scores from Wilcoxon signed-rank tests, comparing predicted footprint scores between control and degron-treated conditions. Let *n* be the number of motif sites, Z-score is computed as

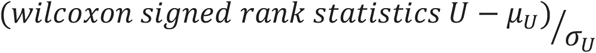

where *µ*_*U*_ = *n*(*n* + 1)/4 and

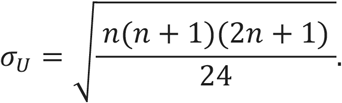

A large Z-score indicates larger differences between two conditions.

### Footprint identifications across different levels of nucleosome depletions

The comparison was performed within the same age group (82 weeks), with weak-depletion DEFND-seq and multiome experiments conducted on the same rat kidney. We restricted our analysis over shared peaks to detect changes across different nucleosome depletion levels. Cell numbers were randomly downsampled to guarantee similar pseudobulk coverage across different conditions. Footprints were identified using TraceBIND with a false discovery rate (FDR) threshold at 5%.

### Footprint identifications during aging in rat kidney proximal tubule cells

Considering that comparison between different age groups might be confounded by changes in cell type composition and overall chromatin accessibility, we restricted our analysis to proximal tubular cells and on shared peaks across all samples. Cell numbers were randomly downsampled to guarantee similar pseudobulk coverage across different samples.

Footprints were identified using TraceBIND with a false discovery rate (FDR) threshold of 5%.

### Standard chromVAR analysis

The positional weight matrices for vertebrates were obtained from the JASPAR2020^61^ database. We then generated a motif matrix using motifmatchr^62^ (p-value cutoff 1e−5) to search for motif sites across all 146,971 peaks on the rat genome assembly GRCr8. Sample-specific chromVAR activity scores were calculated using the RunChromVAR wrapper in Signac^14^ on GRCr8.

### Footprint-refined chromVAR analysis

For each sample, we refined the motif matrix obtained from standard chromVAR^49^ analysis by only retaining TF motif sites overlapping with a TraceBIND-identified sample-specific TF footprint (width <= 120 bps). The resulting refined motif matrix was then used as input to the RunChromVAR wrapper in Signac, with other parameters the same as standard chromVAR.

### Differential chromVAR activity scores between different ages

To assess differential TF activity between age groups, we performed the Wilcoxon rank sum test for each TF, comparing activity levels across cells between different age groups. We reported both the p-value and the normalized Z-score for each comparison. P-values were adjusted for multiple testing. Let *n*_1_ and *n*_2_ be the number of cells in two comparison groups respectively. The Z-score is calculated as

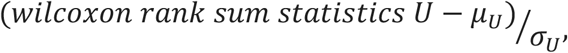

where *µ*_*U*_ = *n*_1_*n*_2_/2 and

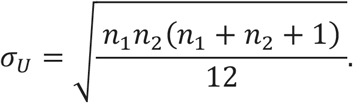

A positive Z-score indicates higher TF activity in older age groups, while a negative Z-score indicates decreased activity with age.

### Splitting peaks into sub-peaks by footprints

Footprints were identified by TraceBIND across all peaks with proximal tubule cells from all age groups. Then the peaks were split up by TraceBIND TF (width <= 120 bps) footprints regions (predicted position ± 3/2 × predicted width) using disjoin from GenomicRanges^64^. Then the subpeak-by-cell count matrix was obtained by FeatureMatrix in Signac. Preprocessing, normalizations, feature selections, dimension reduction and visualizations follow the standard procedures in Signac.

### Epitrace predictions of ages

Age prediction was performed using Epitrace^46^ with default parameters on the rat genome assembly GRCr7 (iterative_time = 1). Inputs included an accessibility matrix derived from scATAC-seq data at both peak and sub-peak levels, along with two sets of clockDMLs: the full set and a subset overlapping TraceBIND-identified TF footprints (width <= 120 bp).

### Overlaps of footprints with Epitrace clockDMLs

The clockDMLs was lifted over from hg38 to GRCr7. Then findoverlaps from GenomicRanges^64^ were performed to find overlaps of TraceBIND-identified footprints and clockDMLs.

## Data availability

The human kidney snATAC datasets were obtained from Wilson et al^22^, Muto et al^23^ and Sheng et al^21^, and are available from the GEO under accession number GSE195460, GSE151302 and GSE115098. The human kidney multiome and snATAC datasets were obtained from Wilson et al^25^ are from GSE232222. Sequencing data for NR3C1, NFAT5 and HIVEP2 CUT&RUN from bulk kidney cortex^22^ are deposited under accession number GSE195443 and GSE220289. The CTCF ChIP-seq from the kidney is deposited under accession number GSM1006886.

## Code Availability

All code used in this study, including the TraceBIND software and the analysis code, can be found on GitHub https://github.com/lyx-lin/TraceBIND.

## Acknowledgement

We thank members of the Zhang and Wilson’s laboratories for useful discussions and critical assessment of this work. We would also like to acknowledge funding support from joint DMS/NIGMS grant DMS-2245575, NIH grants R01GM25301, R01GM149671, and 1R56AG081351, and The Mark Foundation to Y.L. and N.R.Z..

## Author Contribution

Conceptualization, Y.L. and N.R.Z. Algorithm development, Y.L. and N.R.Z. Software implementation, Y.L. Design of computational experiments and data analysis, Y.L., with feedback from W.C.P. and N.R.Z.. Rats DEFND-seq and aging scMultiome experiments, H.W. and W.C.P. Manuscript writing, Y.L. and N.R.Z., with feedback from W.C.P. Supervision, W.C.P. and N.R.Z..

