## Supplementary Figures for "Robust footprinting with sample-specific Tn5 bias correction for bulk and single cell ATAC-seq"

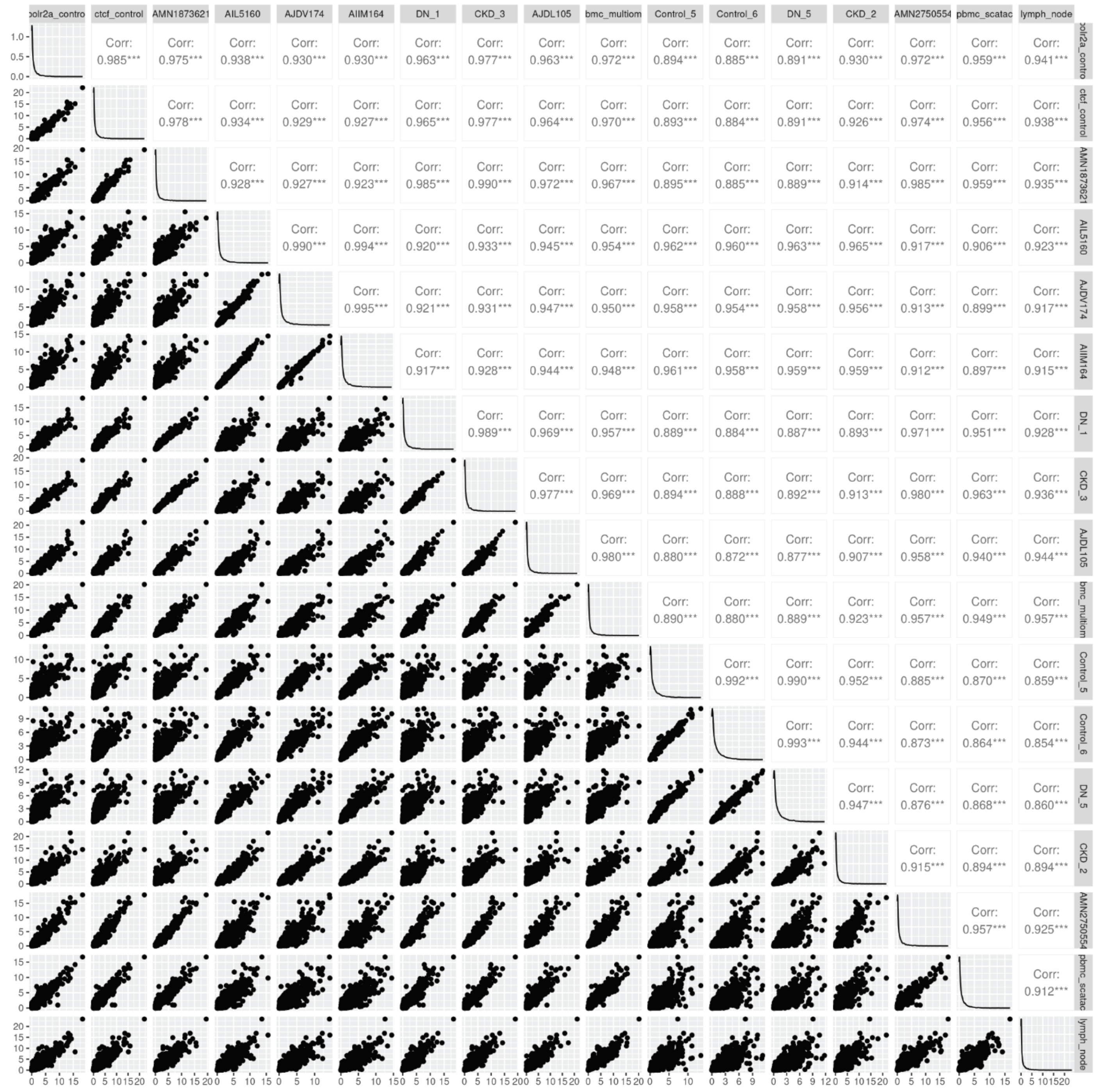


**Suppl Fig1:**
Scatter plots of observed Tn5 bias in mitochondria regions

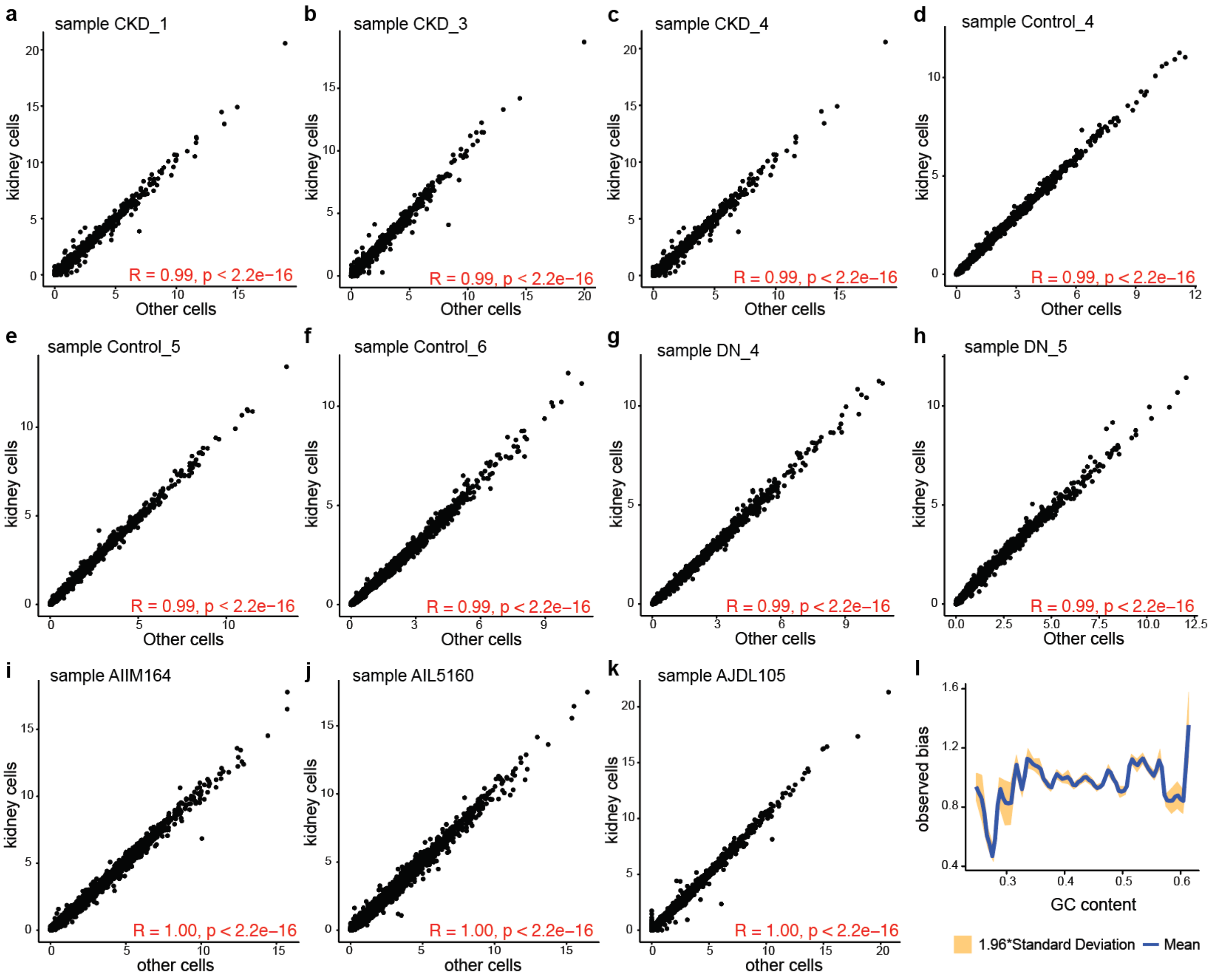


**Suppl Fig2:**

**a-k**: Scatter plots of observed Tn5 bias across different cell types within the same sample. **l**, relationship between GC content and observed bias.

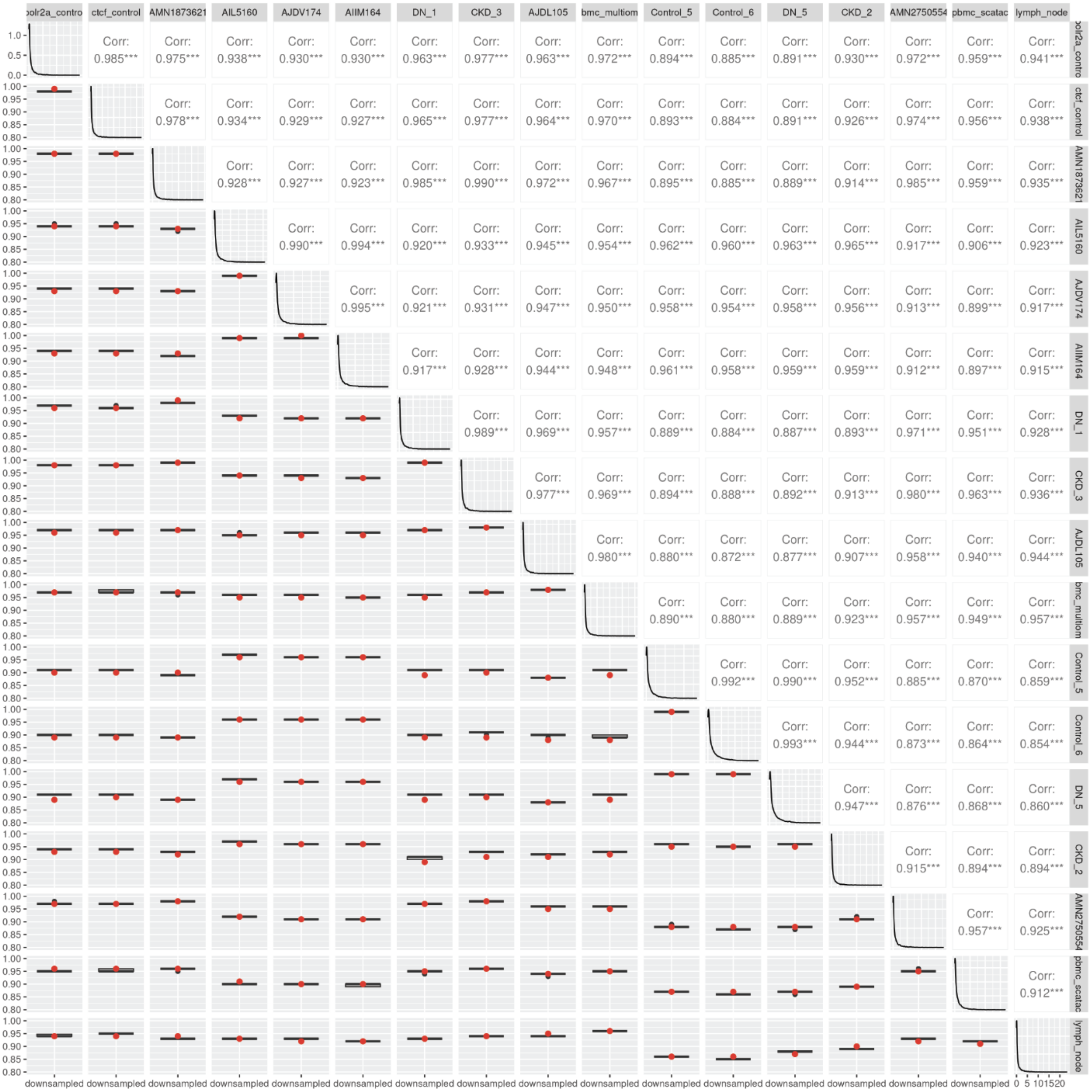


**Suppl Fig3:**

Boxplot shows the distribution of 50 correlation values of observed Tn5 bias after coverage equalization via binomial downsampling. Red dot represents the correlation between samples without downsampling.

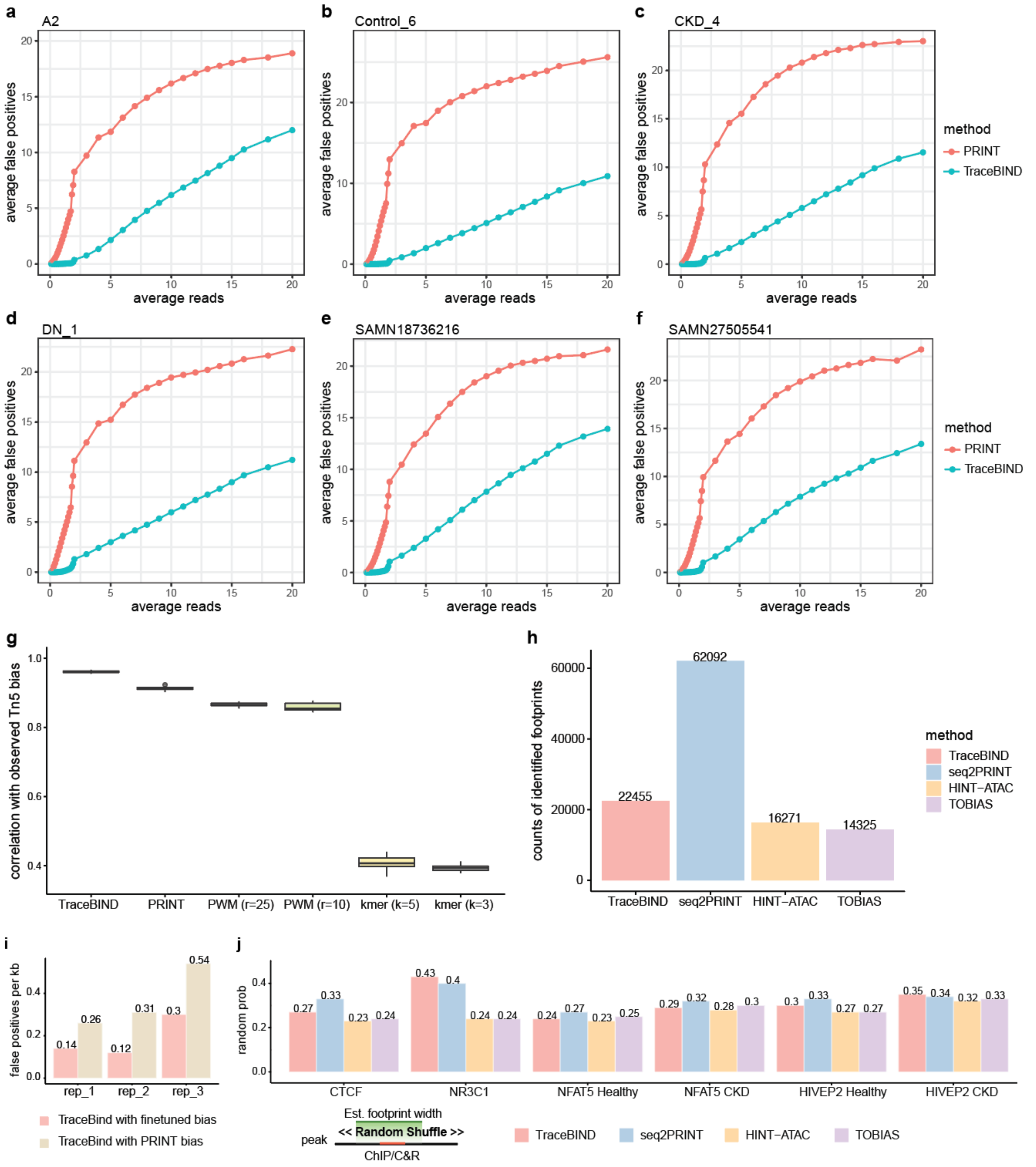


**Suppl Fig4:**

**a-f:** The comparison of average number of false positives in mitochondria test region using different Tn5 bias. **g,** The correlation of predicted and observed Tn5 bias for each method **h,** The counts of identified footprints by different methods with DNase footprints as ground truth. **i,** The false positive rates of TraceBIND with different Tn5 bias. **h,** The random probability by different methods with ChIP/CUT&RUN as ground truth, highlighting the probability of random footprints with reported width overlapping with ChIP/CUT&RUN results by chance.

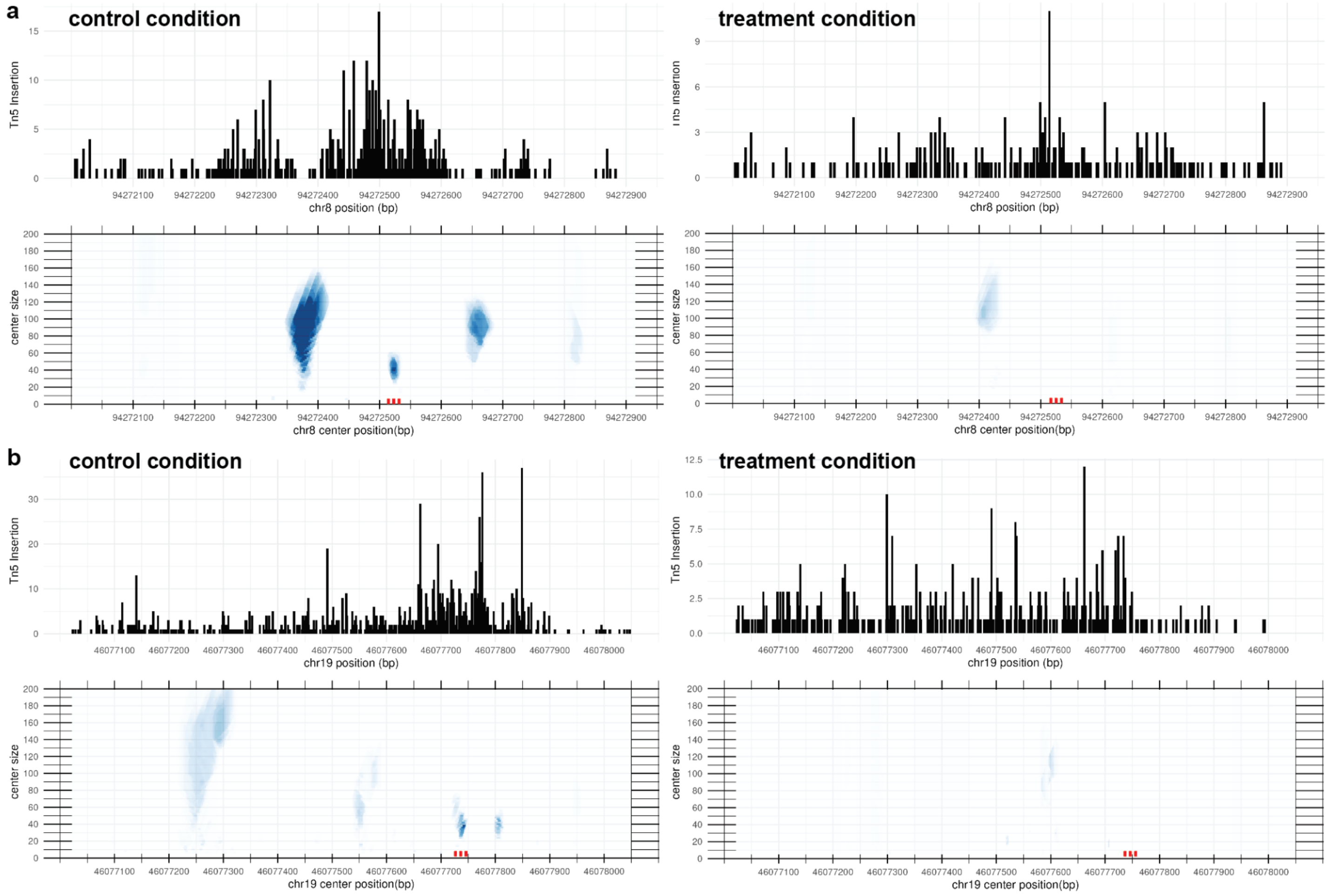


**Suppl Fig5:**

Example regions showing predicted footprints at CTCF motif sites bound in the control but not in the treatment condition. Red dotted lines indicate CTCF motif positions.

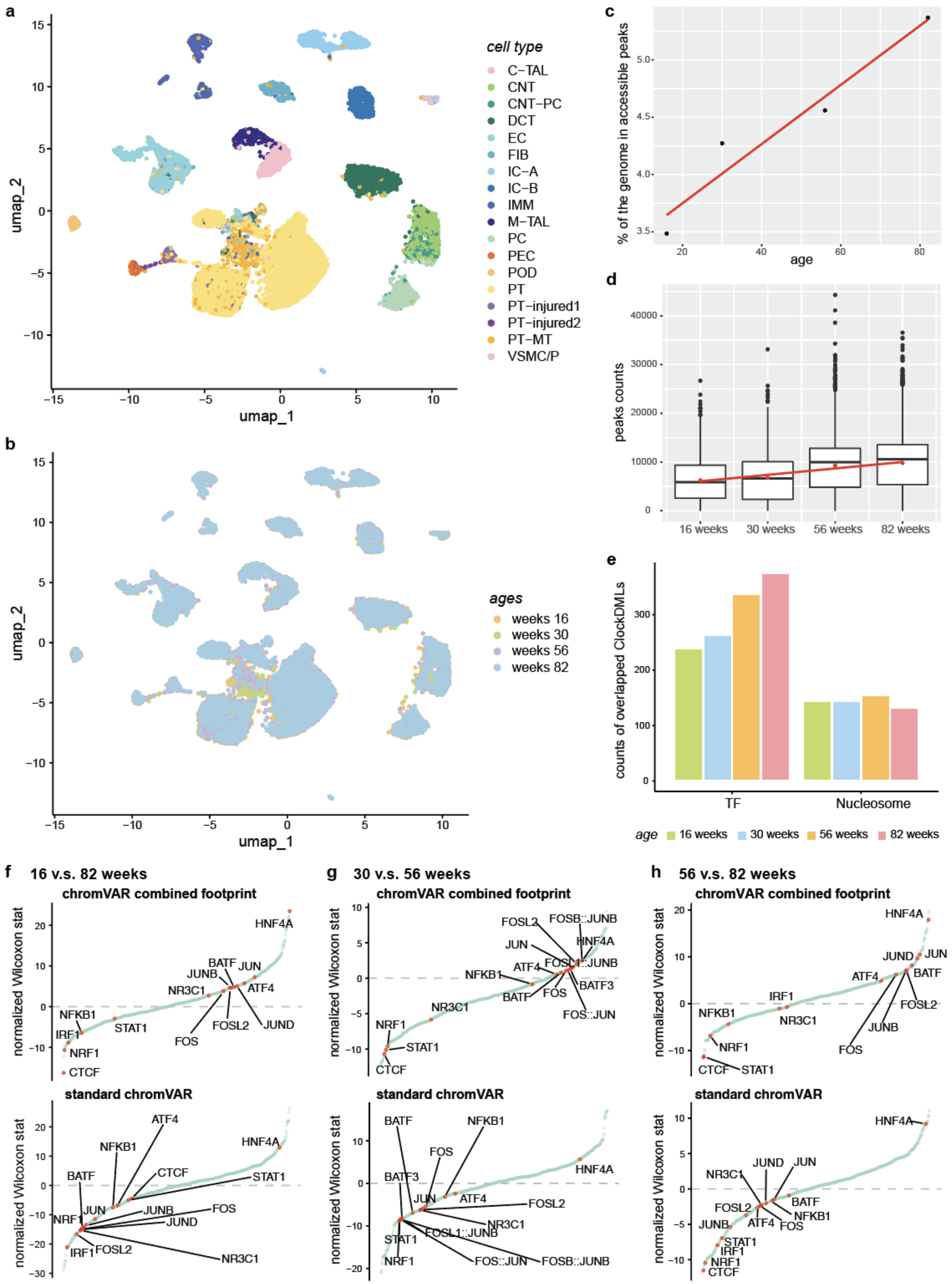
a
cell type
C−TAL
CNT
CNT−PC
DCT
EC
FIB
IC−A
IC−B
IMM
M−TAL
PC
PEC
POD
PT
PT−injured1
PT−injured2
PT−MT
VSMC/P
umap_1
umap_2
b
ages
weeks 16
weeks 30
weeks 56
weeks 82
c
% of the genome in accessible peaks
age
d
peaks counts
16 weeks
30 weeks
56 weeks
82 weeks
e
counts of overlapped ClockDMLs
TF
Nucleosome
age
16 weeks
30 weeks
56 weeks
82 weeks
f
16 v.s. 82 weeks
g
30 v.s. 56 weeks
h
56 v.s. 82 weeks
chromVAR combined footprint
standard chromVAR
normalized Wilcoxon stat
HNF4A
JUN
JUND
JUNB
BATF
BATF3
ATF4
FOS
FOSL2
FOSL1::JUNB
FOSB::JUNB
FOS::JUN
NR3C1
NFKB1
IRF1
NRF1
STAT1
CTCF

**a, b**: visualizations of scATAC data based on peak level colored by cell types and samples (after integration). **c, d**, percentage of genome covered by accessible regions (peaks) and the number of peaks per cell across different age groups. **e**, numbers of footprints overlapping with age-associated hypomethylation sites. **f**, Comparison of TF activity scores across different age groups using standard chromVAR and footprint-informed chromVAR. Positive normalized Wilcoxon statistics means higher activities in older groups.

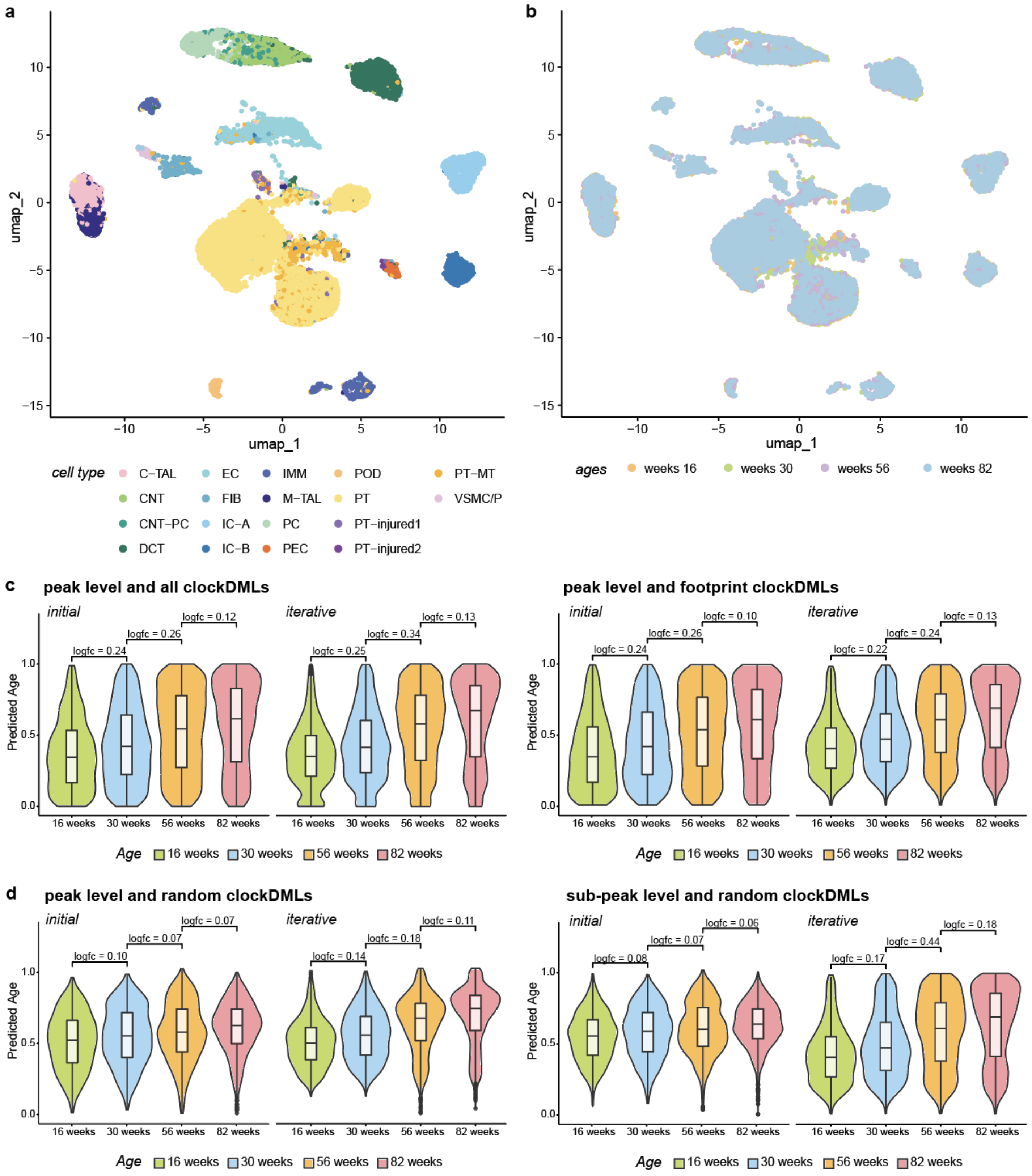


**Suppl Fig 7.**

**a, b**: visualizations of scATAC data based on sub-peak level split by TraceBIND footprints, colored by cell types and samples (after integration). **c, d**: Epitrace results using different sets of ClockDMLs and peak resolution levels.
